# Identification of novel kolmioviruses in diverse invertebrates

**DOI:** 10.64898/2026.09.02.748743

**Authors:** Kyoka Miyata, Mai Kishimoto, Hana Sasaki, Masayuki Horie

## Abstract

Kolmioviruses (KoVs; family *Kolmioviridae*) are circular, single-stranded RNA viruses that have recently been discovered in a wide range of animal species. However, invertebrate-associated KoVs have so far been identified only in termites, and their diversity and evolution remain largely unexplored. Here, we comprehensively analyzed publicly available animal RNA-seq datasets and identified six novel KoVs associated with one fish and five invertebrate species: coho salmon (*Oncorhynchus kisutch* [Walbaum, 1792]), spotted babylon (*Babylonia areolata* [Link, 1807]), Atlantic pearl oyster (*Pinctada imbricata* [Röding, 1798]), small emerald moth (*Hemistola chrysoprasaria* [Esper, 1795]), codling moth (*Cydia pomonella* [Linnaeus, 1758]), and cocoa pod borer (*Conopomorpha cramerella* [Snellen, 1904]).

---

Kolmioviruses (KoVs; family *Kolmioviridae*) are small, circular, and single-stranded RNA viruses with highly self-complementary genomes (1). All known kolmioviruses contain an antigenomic-sense ORF encoding the delta antigen (DAg), typically as their sole protein-coding ORF, whereas canary kolmiovirus additionally contains a genomic-sense ORF (ORF2) (2). For several decades after the discovery of hepatitis delta virus (HDV) in the 1970s (3), no related viruses were identified. Since 2018, however, KoVs have been detected in diverse animal species, expanding our knowledge of host range (4–10). Nevertheless, invertebrate-associated KoVs have so far been reported only in termites (6, 11). Here, we surveyed publicly available animal RNA-seq datasets for novel KoVs, focusing on the largely unexplored diversity of invertebrate-associated KoVs.

We screened 500,276 vertebrate- and 1,049,723 invertebrate-derived RNA-seq datasets in the NCBI Sequence Read Archive (SRA) (12) using SRAminer (13). For KoV-positive datasets, raw reads were preprocessed with fastp (14) and mapped to the corresponding host genomes using HISAT2 (15). Unmapped reads were then extracted with SAMtools (16) and *de novo* assembled using rnaviralSPAdes (17) and Trinity (18). When multiple KoV-positive datasets were available for the same host species, their unmapped reads were co-assembled. Resulting contigs were screened by BLASTx searches (19) against known KoV DAg sequences, followed by BLASTx and BLASTp searches against the clustered nr database (12). For contigs showing sequence similarity to KoV DAg, reads from each RNA-seq dataset were individually mapped back to the corresponding contigs, and regions with a read depth of <2 were trimmed.

We identified six novel KoVs, including one vertebrate-associated KoV from coho salmon (*Oncorhynchus kisutch* [Walbaum, 1792]) and five invertebrate-associated KoVs from two molluscan hosts, spotted babylon (*Babylonia areolata* [Link, 1807]) and Atlantic pearl oyster (*Pinctada imbricata* [Röding, 1798]), and three lepidopteran hosts, small emerald moth

(*Hemistola chrysoprasaria* [Esper, 1795]), codling moth (*Cydia pomonella* [Linnaeus, 1758]), and cocoa pod borer (*Conopomorpha cramerella* [Snellen, 1904]) (Table 1). Complete genome sequences were obtained for B. areolata KoV (baKoV) and H. chrysoprasaria KoV (hcKoV), whereas only partial sequences were recovered for the remaining KoVs (Fig. 1A).

**Table 1.**
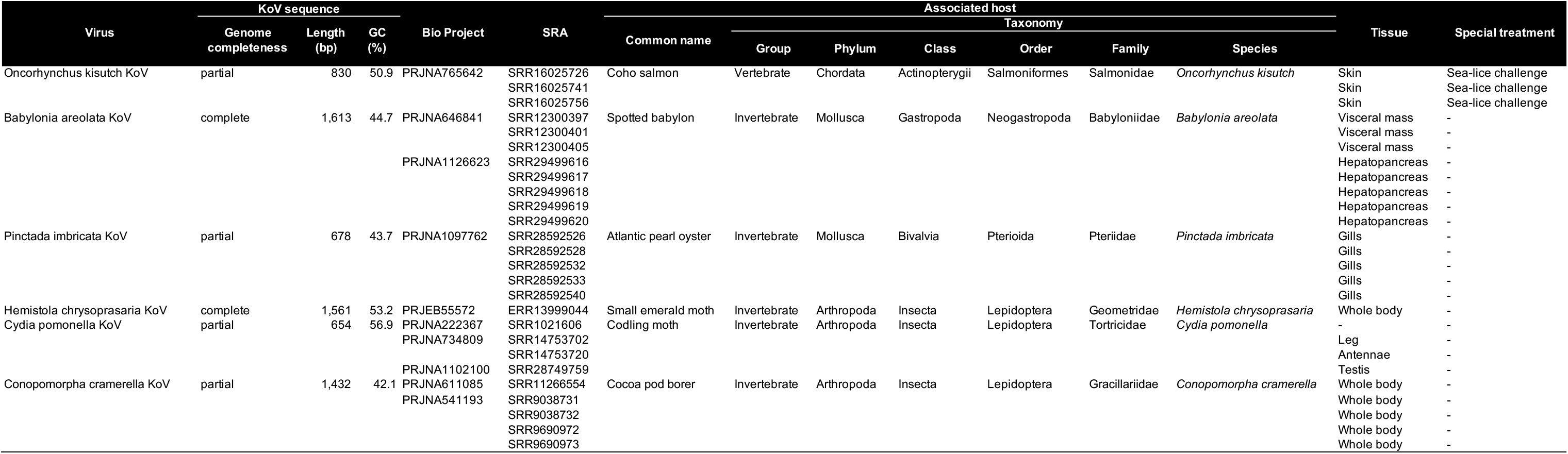
Kolmiovirus sequences identified in this study.

**Fig. 1.**
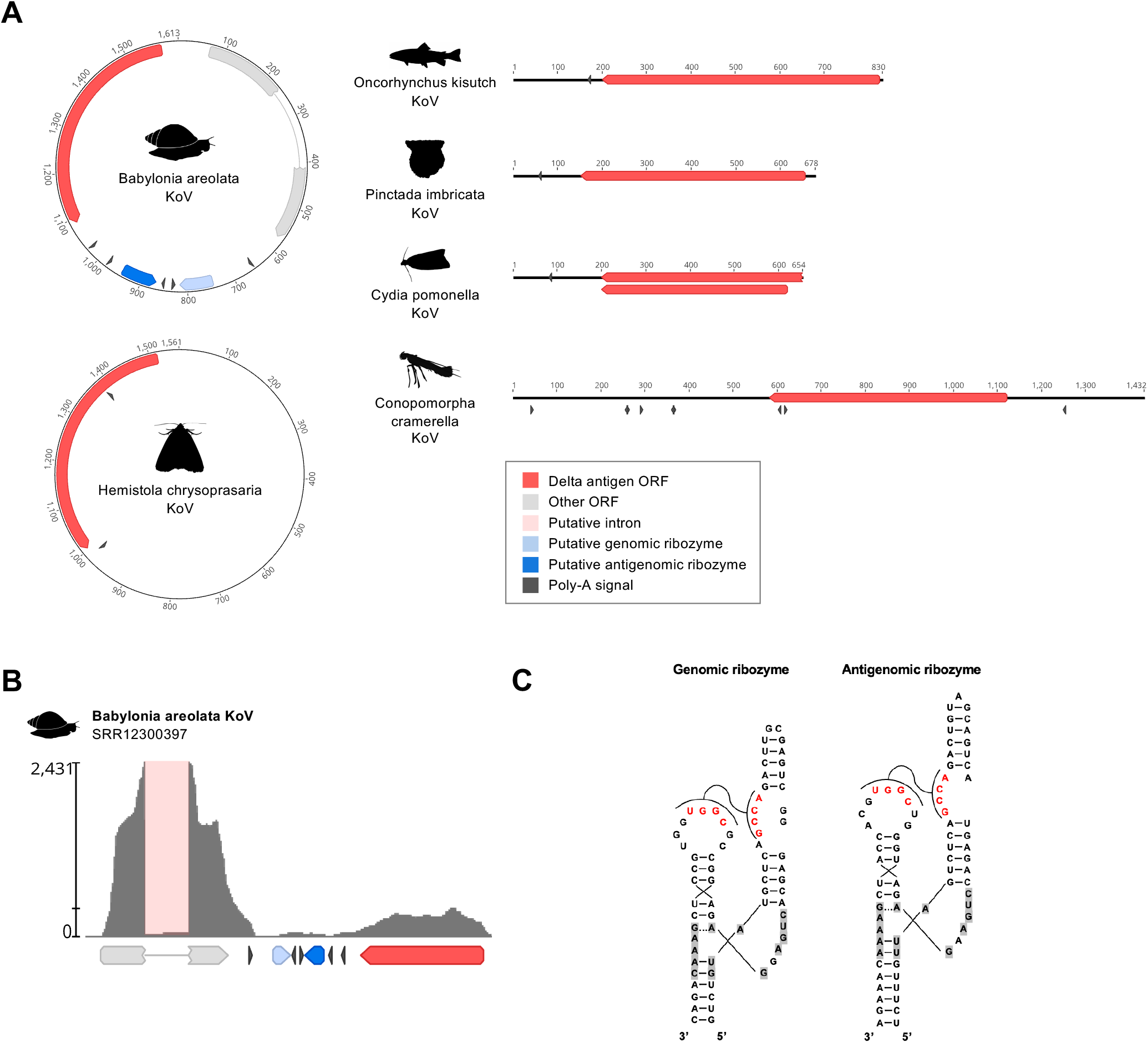
Identification and characterization of novel KoVs. (**A**) Genome organization of novel KoVs identified in this study. Colored arrows indicate annotations (ORFs, ribozymes, and poly-A signals). The numbers indicate nucleotide positions. Animal silhouettes were obtained from PhyloPic (https://www.phylopic.org/). (**B**) Mapping coverage of original short reads of Babylonia areolata KoV. Colored arrows indicate annotations (ORFs, ribozymes, poly-A signals). Light pink box indicates putative intron. (**C**) Predicted genomic and antigenomic ribozyme structures of Babylonia areolata KoV. The most frequent nucleotides in the catalytic core are highlighted in gray. Conserved loop-loop interactions are indicated in red.

We next characterized the genome structure of baKoV. Read-mapping analysis revealed two distinct coverage peaks, consistent with an ambisense transcription pattern (2). Analysis of the mapped reads also identified alternative splicing of the transcript (Fig. 1B). Although the unspliced transcript contained only a short ORF, the spliced transcript contained a longer putative ORF. Prediction of self-cleaving ribozymes revealed type III hammerhead ribozymes in both the genomic and antigenomic strands (Fig. 1C).

Phylogenetic analysis based on DAg amino acid sequences revealed two major lineages of invertebrate-associated KoVs (Fig. 2). The Mollusca lineage comprised the newly identified Mollusca-associated KoVs, whereas the Arthropoda-associated lineage comprised the newly identified Lepidoptera-associated KoVs together with previously reported termite KoVs. Chusan Island toad virus 1 also clustered within the Arthropoda-associated lineage, consistent with our detection of an identical KoV sequence in a Lepidoptera RNA-seq dataset (SRR3400941).

**Fig. 2.**
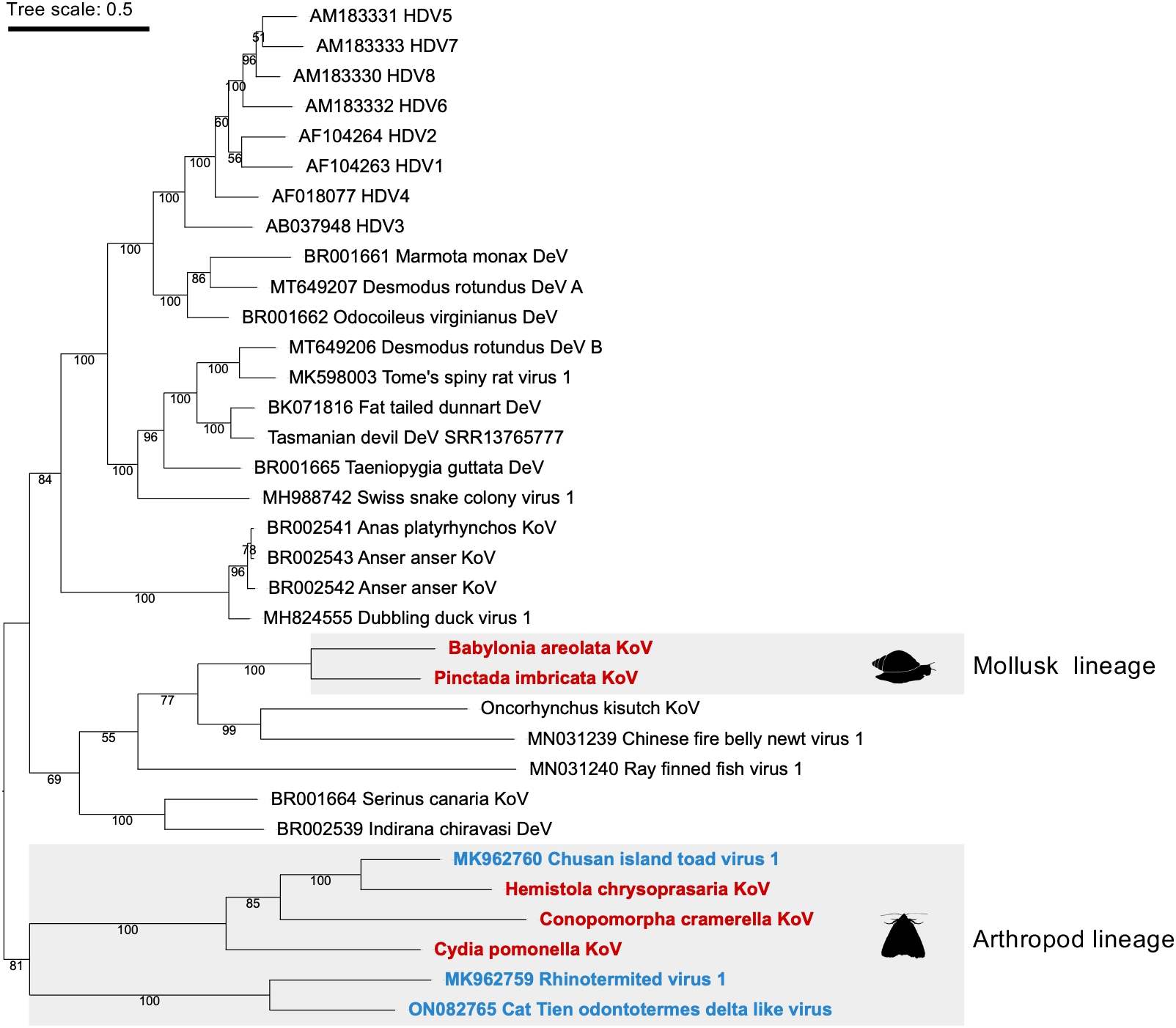
Phylogenetic relationship between vertebrate and invertebrate KoVs. **(A)** Phylogenetic tree was inferred by the Bayesian Markov chain Monte Carlo method based on an amino acid sequence alignment of DAg. Posterior probabilities are shown for all nodes. Invertebrate-associated KoVs identified in this study are shown in red, and previously reported invertebrate-associated KoVs are shown in blue. The scale bar indicates the number of amino acid substitutions per site. Animal silhouettes representing each host group were obtained from PhyloPic (https://www.phylopic.org/).

In this study, we identified KoVs from non-termite invertebrates for the first time, substantially expanding the known host range of KoVs. Phylogenetic analysis revealed that invertebrate-associated KoVs form two major lineages, corresponding to Arthropoda- and Mollusca-associated KoVs, suggesting diversification across distinct invertebrate phyla. Detailed characterization of baKoV provided evidence for an ambisense genome organization, alternative splicing, and the presence of type III hammerhead ribozymes in both the genomic and antigenomic strands. Together, these findings broaden our understanding of the diversity and evolution of KoVs, particularly in invertebrates, and suggest that complex genome organization and gene expression strategies may be more widespread within the family than previously recognized.

## Acknowledgements

This study was supported by KAKENHI grant numbers 21H01199 (MH), 22K19234 (MH), 23K20902 (MH), 24K21922 (MH), and 24K18455 (MK) and the 2024/2025/2026 Osaka Metropolitan University (OMU) Strategic Research Promotion Project (Young Researcher) (MK).

